# Pancreatic Islets Communicate With the Brain *via* Vagal Sensory Neurons

**DOI:** 10.1101/780395

**Authors:** Madina Makhmutova, Jonathan Weitz, Alejandro Tamayo, Elizabeth Pereira, Joana Almaça, Rayner Rodriguez-Diaz, Alejandro Caicedo

## Abstract

Depleting visceral sensory nerves affects pancreatic islet function, glucose metabolism and diabetes onset, but how islet endocrine cells interact with sensory neurons has not been studied. Here we show that the pancreatic islet is innervated by vagal sensory axons expressing substance P, calcitonin-gene related peptide, and serotonin receptor 5HT3R. Vagal neurons projecting to the pancreas terminate in the commissural nucleus of the solitary tract. These neurons respond to chemical but not mechanical stimulation of the pancreas. By recording activity from nodose neurons *in vivo* and from sensory axons in living pancreas slices, we show that sensory nerves respond to serotonin secreted from stimulated beta cells. Serotonin is co-released with insulin and therefore conveys information about the secretory state of beta cells *via* vagal afferent nerves. Our study thus establishes that pancreatic islets communicate with the brain using the neural route and identifies serotonin signaling as a peripheral transduction mechanism.

## INTRODUCTION

Recent neurobiological studies are revealing how the brain communicates with visceral organs. This includes the pancreatic islet of Langerhans, a mini-organ whose endocrine cells secrete insulin and glucagon, two hormones that are crucial for glucose homeostasis. At their discovery in the late 19th century, Paul Langerhans described that pancreatic islets are richly innervated (Langerhans and Morrison, 1937). Claude Bernard had reported earlier that puncturing the floor of the fourth ventricle induces diabetes (Bernard, 1849). Since then, it is generally assumed that the brain helps control glucose homeostasis, presumably *via* autonomic nerves that regulate pancreatic endocrine function. It is not until recently, however, that detailed neuroanatomical studies revealed how the efferent branches of the autonomic nervous system, the sympathetic and parasympathetic nerves, innervate distinct cell populations within the islet (Almaça et al., 2018; Rodriguez-Diaz et al., 2011; Taborsky, 2011). By contrast, little is known about the sensory innervation of the islet. Because visceral sensory innervation is a crucial component of homeostatic regulatory circuits (Chang et al., 2015; Li & Owyang, 1993; Nonomura et al., 2016; Williams et al., 2016), there is a need to understand how islets signal to sensory fibers.

A wide body of literature shows that chemical and genetic sensory denervation affects physiologic and pathophysiologic processes in the pancreas, including insulin secretion and diabetes onset (Bou Karam et al., 2018; Gram et al., 2007; Ikeura et al., 2007; Karlsson et al., 1992; Li et al., 2013; Liddle, 2007; Motter & Ahern, 2008; Noble et al., 2006; Razavi et al., 2006; Riera et al., 2014; Warzecha et al., 2001). Little is known, however, about the basic patterns of pancreatic sensory innervation and its transduction mechanisms. Tracing studies showed that vagal afferent axons innervate the pancreas (Berthoud & Neuhuber, 2000; Carobi, 1987; Fasanella et al., 2008; Neuhuber, 1989; Sharkey & Williams, 1983). In rodents, sensory fibers are localized preferentially to the periphery of the islet forming a dense superficial network (Gram et al., 2007; Hannibal & Fahrenkrug, 2000; Karlsson et al., 1992; Lindsay et al., 2005; Pettersson et al., 1986; Seifert et al., 1985; Sternini et al., 1992). Functional studies found that dispersed sensory afferent neurons of the vagal nodose ganglion respond to pancreatic and gastrointestinal stimuli, as well as to islet-derived stimuli (Ayush et al., 2015; Iwasaki et al. 2017; Iwasaki et al., 2013; Peters et al., 2004; Simasko & Ritter, 2003). These findings suggest that the vagus nerve could sense the islet microenvironment, yet this notion has not been explored. We still don’t know what activates the vagal islet-brain axis and how changes in the islet microenvironment affect vagus activity.

Serotonin is one of the strongest stimuli for vagal afferent neurons (Bellono et al., 2017; Bornstein et al., 2015; Hillsley & Grundy, 1998; Y. Li et al., 2001). Within the pancreas, the insulin-producing beta cell is likely the sole source of serotonin. Beta cells secrete serotonin to communicate with neighboring cells (Almaça et al., 2016; El-Merahbi et al., 2015; Goyvaerts et al., 2016; K. Kim et al., 2015; Kim et al., 2017; Ohara-Imaizumi et al., 2013; Ohta et al., 2011; Teitelman et al., 1987). We therefore hypothesized that pancreatic islets use serotonin as a signaling molecule to communicate with the brain *via* vagal afferents. We used anatomical and physiological tools to characterize the sensory innervation patterns in the islet, to identify brain regions that receive pancreatic input, to examine neuronal response profiles in the nodose ganglion during pancreatic manipulation *in vivo*, and to demonstrate that vagal sensory neurons innervating the islet respond to serotonin released from beta cells.

## RESULTS

### The mouse pancreas is innervated by vagal afferent neurons that project to the commissural nucleus of the solitary tract

We investigated the anatomical components of the vagal pancreas-brain axis (Figure 1A) using immunohistochemistry and neuronal tracing. To examine the pattern of sensory innervation of the mouse and human pancreas we used immunohistochemistry for the common sensory neuronal markers substance P and calcitonin-gene related peptide (CGRP; Figures 1B-1E). Islets were innervated by substance P and CGRP immunoreactive axons. The density of innervation within the islet was significantly higher than in the surrounding exocrine tissue (Figures 1B and 1C). CGRP-labeled axons did not express choline acetyltransferase (ChAT), a marker for parasympathetic efferent terminals (Figures 1D and 1D’). We did not observe any gender or strain differences within the analyzed mouse strains (129, BALB6-C, C57Bl6; Figure S1). However, males of the non-obese diabetic (NOD) mouse strain had a significantly lower density of sensory fibers within the islet (Figure S1). Human pancreatic islets were similarly innervated by substance P immunoreactive axons (Figures 1E and 1E’).

**Figure 1.**
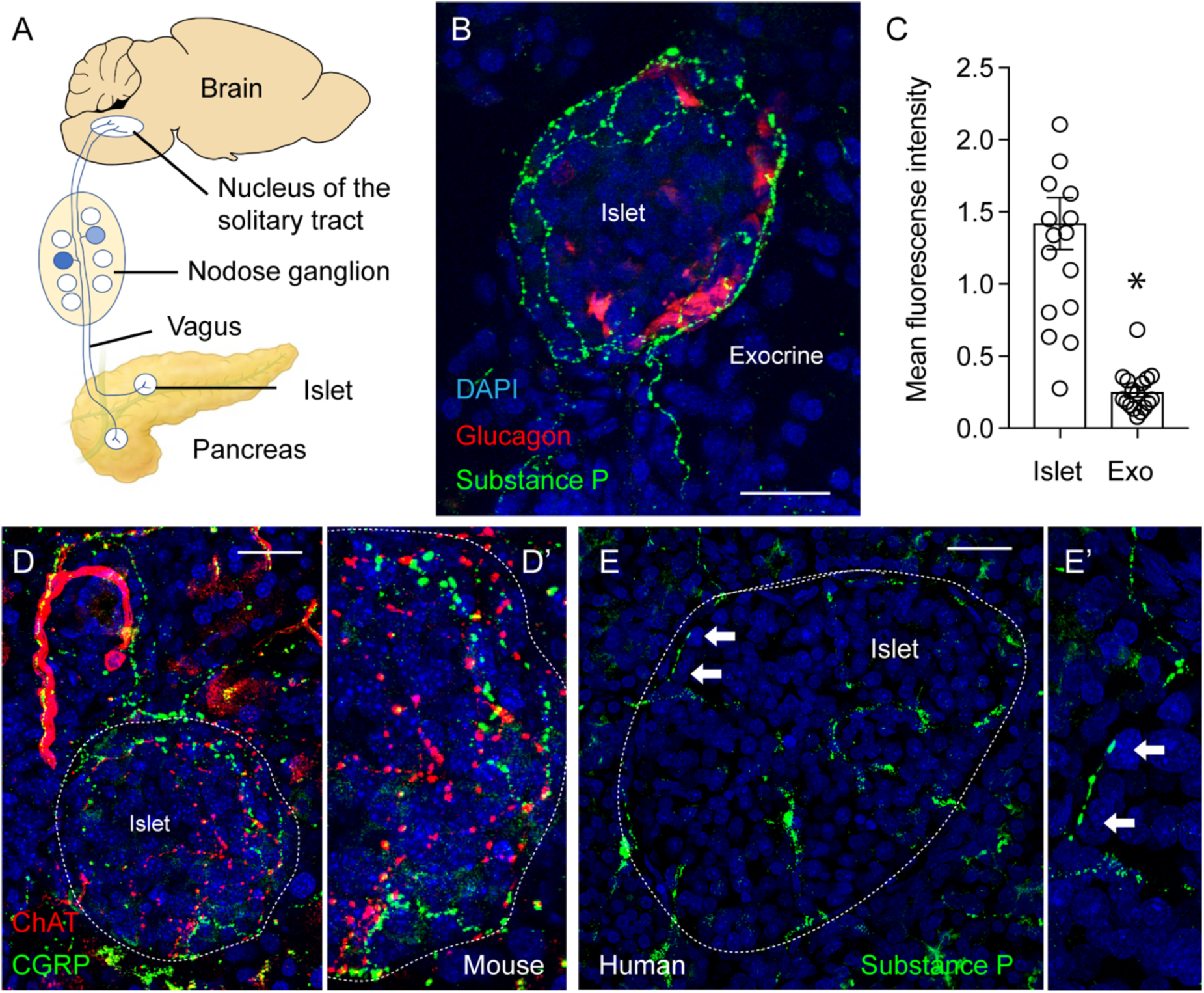
Mouse and human pancreatic islets are innervated by SP- and CGRP-positive sensory fibers. (A) Cartoon representation of the vagal islet-brain axis. (B) Mouse pancreas section showing the innervation pattern of substance P-positive sensory axons (green) in an islet. Endocrine alpha cells labeled for glucagon are shown in red. Scale bar, 20 μM. All immunostaining figures are shown as z-stacks of confocal images. (C) Quantification of the mean fluorescence intensity of substance P immunostaining in the islet and the surrounding exocrine tissue (n = 18 mice, 3-7 islets per animal; mean +/- SEM, paired t-test; for data showing innervation densities in mice of different genders and strains, see Figure S1). (D-D’) Innervation pattern of CGRP-positive sensory axons (green) and cholinergic parasympathetic axons (red) in an islet of a ChAT-GFP reporter mouse. Scale bar, 20 μm. (E-E’) Innervation pattern of substance P-positive sensory axons in a human pancreatic islet (green). Arrows point at axon varicosities at the border of the islet. Scale bar, 20 μm.

To identify vagal sensory neurons projecting to the pancreas and examine their terminal fields in the brain, we performed retrograde tracing by infusing the pancreas with tracers through the common bile duct, as previously described (Guo et al., 2014; Figure 2). We compared tracing efficiencies of three tracers: fast blue (Figures 2A and 2B), cholera toxin subunit beta (CTB, Figure 2C), and retrograde AAV-hSyn-mCherry (Figures 2D and 2E). We obtained similar results with all three tracers at the level of the nodose ganglion (the vagal sensory ganglion), with CTB tracing showing a lower tracing efficiency (Figures 2B and 2C). Retrograde tracing from the pancreas to the nodose ganglion using CTB labeled 72 +/- 9 neurons in the right ganglion and 100 +/- 8 neurons in the left ganglion (n = 3 mice). In line with previous observations we could not find strict topographic organization of these neurons (Fasanella et al., 2008; Zhuo et al., 1997). We estimated that from 2–6% of nodose neurons were traced from the pancreas. By contrast, 50% of nodose ganglion neurons were traced when CTB was injected intraperitoneally (Figure 2C). Neurons in the trigeminal ganglion could not be traced from the pancreas (negative control, Figure 2C).

**Figure 2.**
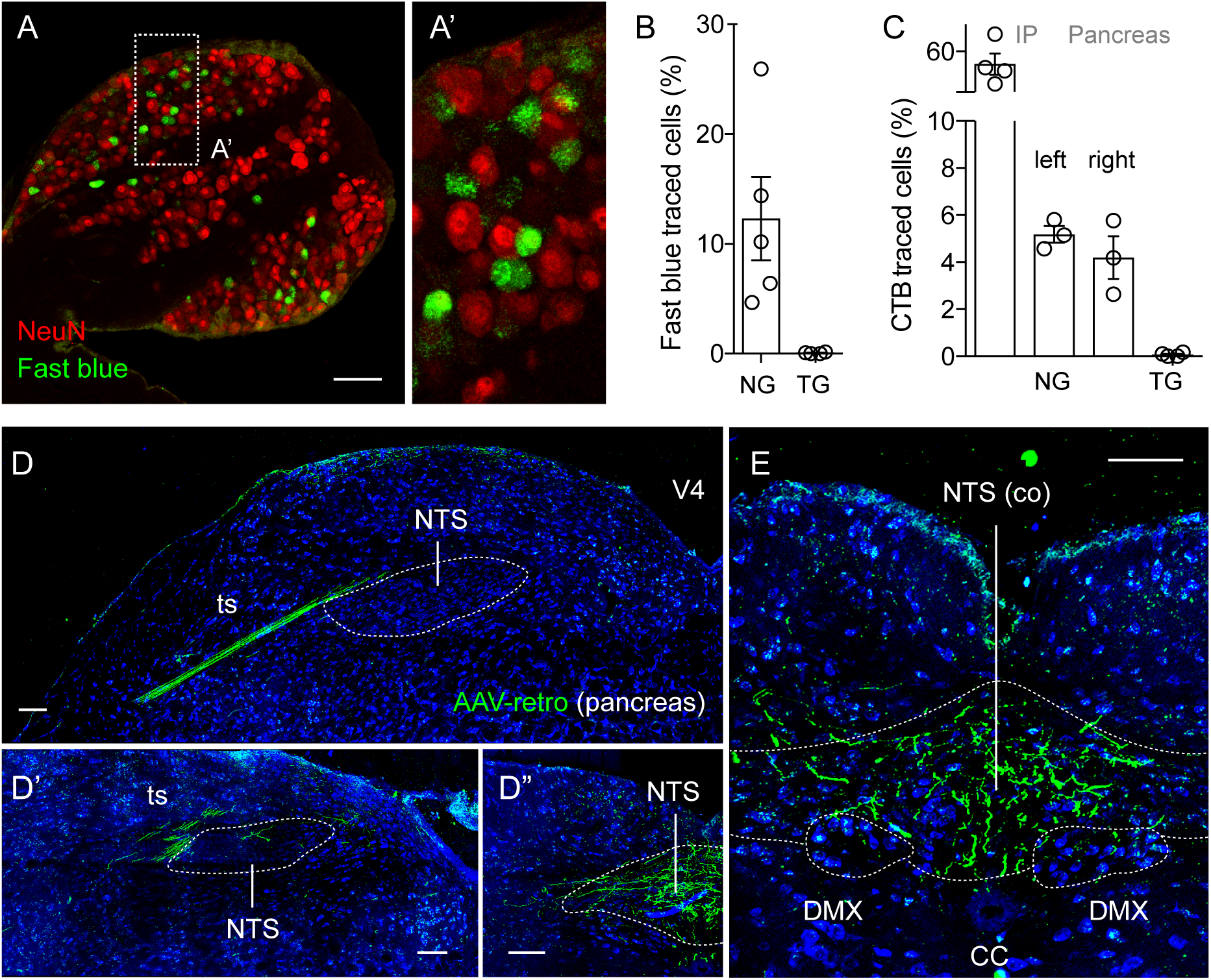
Pancreatic vagal afferents project to the commissural region of the nucleus of the solitary tract (NTS) (A-A’) Section of a mouse nodose ganglion showing retrogradely traced neurons (green) 1 week after intraductal pancreas infusion of the neuronal tracer fast blue. NeuN staining (red) visualizes all neurons. (B) Quantification of data as in A showing the percentage of neurons that is traced with Fast Blue in the left nodose ganglion and trigeminal ganglion (negative control); (n = 5 mice, 5-9 sections per ganglion). (C) Quantification of the percentage of neurons traced with cholera toxin B (CTB) in the left nodose ganglion after IP injection of the tracer (positive control) and the left and right nodose ganglia and trigeminal ganglion (negative control) after intraductal pancreas infusion of CTB (n = 3 mice, 5-9 sections per ganglion). (D-D”) Sequential images of mouse brainstem sections (rostral to caudal) 4 weeks after intraductal pancreas infusion of the neuronal tracer AAV-retro-hSyn-mCherry. Traced fibers are shown in green. Cytoarchitecture of the brainstem is identified with Nissl stain (blue). (E) Section at the level of the caudal commissural NTS containing the final terminal field of neurons traced from the pancreas. Scale bars in all panels, 100 μm.

Although fast blue and CTB-traced neurons were seen in the nodose ganglion, we could not identify the terminal fields of these neurons in the brain. The retrograde AAV-hSyn-mCherry tracer, by contrast, worked efficiently to trace the whole pathway from the pancreas to the brain (Figures 2D and 2E). We found that the central projections of pancreas-innervating vagal sensory neurons terminated in the commissural regions of the caudal nucleus of the solitary tract.

We next sought to identify projections of vagal sensory neurons in the pancreas. We performed anterograde tracing by injecting AAV-CMV-mCherry into the nodose ganglion, as described (Chang et al., 2015). Four weeks after tracer injection, sensory fibers could be traced to the pancreas, thus confirming previous findings (Neuhuber, 1989). Traced fibers were also seen in the brain and duodenum, matching the pattern previously reported (Chang et al., 2015; Figure S2).

### *In vivo* imaging of neuronal activity in the intact nodose ganglion in response to pancreas-specific stimulation

To characterize sensory innervation of the pancreas physiologically, we adapted a technique for *in vivo* imaging of the nodose ganglion (Williams et al., 2016). We developed a surgical approach that allowed imaging of cytoplasmic Ca^2+^ of neurons in the intact nodose ganglion without cutting the central branch of the vagus nerve (Figures 3A-3C, Movie S1). Using this method, we not only recorded stimulus-induced neuronal responses, but also observed that most vagal sensory neurons had a baseline activity that disappeared after cutting the central branch.

**Figure 3.**
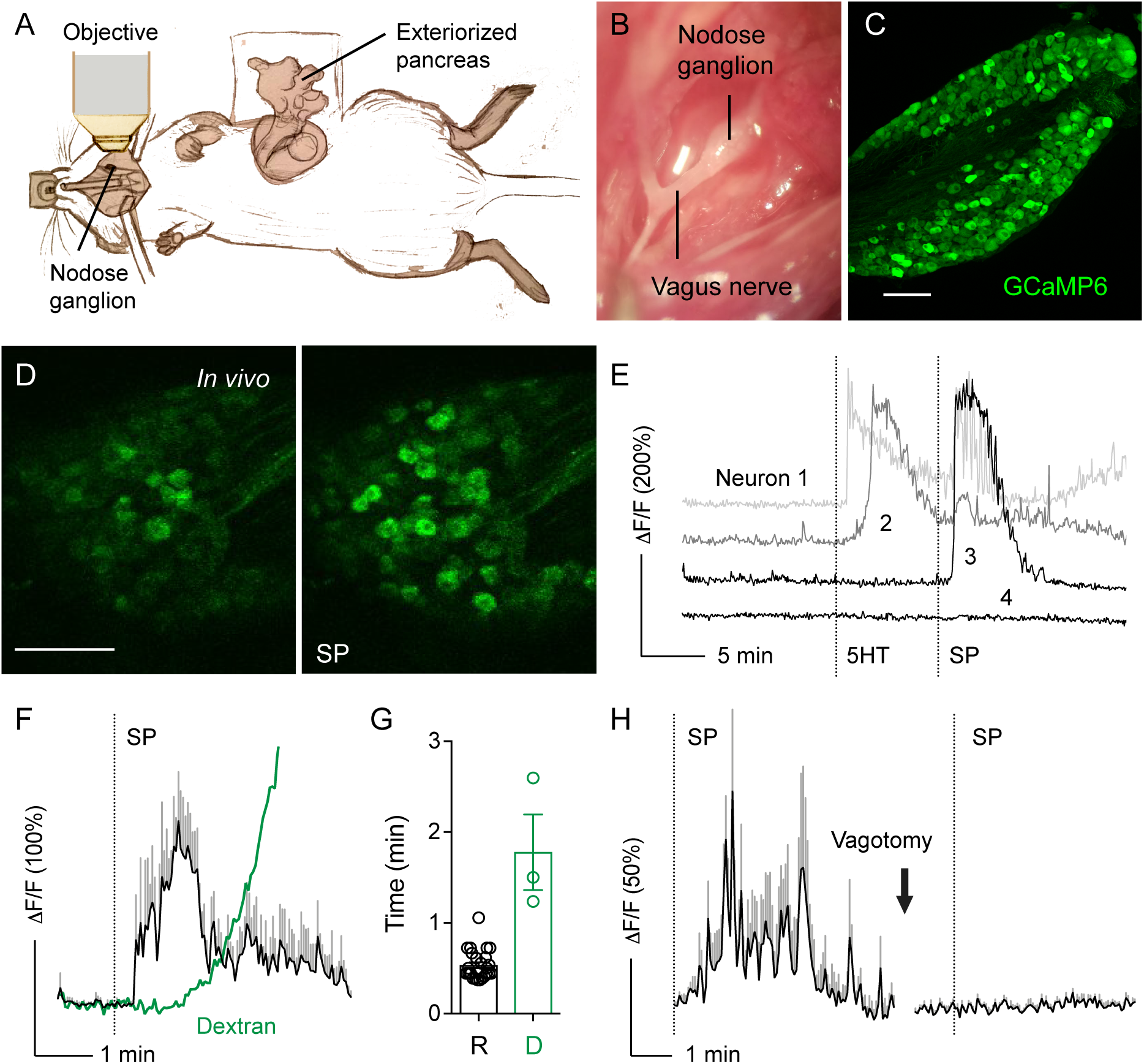
*In vivo* imaging of the intact nodose ganglion in response to pancreas-specific stimulation. (A) Cartoon illustrating the experimental setup, where the intact nodose ganglion is exposed for Ca^2+^ imaging under a confocal microscope and the pancreas is exteriorized for topical stimulus application. See Methods for detailed description of surgeries. (B) Microphotograph of the exposed nodose ganglion. (C) Confocal image of the nodose ganglion from a Pirt-GCaMP6 mouse, which expresses the Ca^2+^ indicator GCaMP6 in all nodose ganglion neurons. (D) Sequential images of nodose ganglion neurons displaying Ca^2+^responses (changes in GCaMP6 fluorescence, green) to topical stimulation of the pancreas with substance P (SP). Scale bars in C and D, 100 μm. (E) Representative traces of neurons recorded in D responding to 5HT (1 mM), SP (100 μM), or both. (F) Traces comparing the time course of neuronal Ca^2+^ responses to substance P (SP, 100 μM, black, average +/- SEM of 5 neurons) with that of the diffusion of fluorescent dextran (3 kDa) into the blood stream (green). A mixture of both substances was applied topically to the exteriorized pancreas and their fluorescence intensities measured in the nodose ganglion. (G) Quantification of data as in F showing the latencies of the Ca^2+^ response to substance P (R) and dextran diffusion into blood vessels of the nodose ganglion (D). Latency of fluorescence appearance was calculated as the first time point after topical application at which the mean fluorescence intensity was at least 3 SD above baseline fluorescence fluctuation (n = 3 animals). (H) Vagotomy abolished Ca^2+^ responses to substance P (SP). Trace shows an average of the same 10 neurons before and after vagotomy (representative of 3 animals).

Specific sensory neuronal markers for the population of vagal neurons innervating the pancreas have not been identified yet. Thus, we recorded activity from all nodose neurons using mice expressing the Cre-dependent Ca^2+^ indicator GCaMP6 driven by the general sensory neuronal promoters Pirt or Snap25 (Madisen et al., 2015; Shin et al., 2014; Figure 3C). There was no difference in Ca^2+^ responses between the two strains. To stimulate the pancreas specifically and to control for non-specific responses induced outside of the pancreas, we infused stimuli either through the common bile duct (intraductally) or applied them topically to the exteriorized pancreas (Figure 3A). Intraductal administration of stimuli accesses the pancreas “inside out”. This procedure allows the stimuli to fill the pancreatic ductal system and diffuse into the periductal space, where the majority of the pancreatic islets are located. Topical stimulus administration, by contrast, accesses the pancreas “outside in” and primarily diffuses into exocrine tissues. Using this approach, responses could be clearly detected in the nodose ganglion neurons, with a strong signal to noise ratio (Figures 3D and 3E).

To control for systemic distribution of topically applied substances, we applied a mixture of a fluorescent dextran (3 kDa) together with a chemical stimulus (substance P, 100 μM) to the exteriorized pancreas. We recorded neuronal Ca^2+^ responses and the appearance of the dextran fluorescence in the ganglion (Figures 3F and 3G). The average latency of the neuronal response was smaller than that of the appearance of a dextran in the circulation (32 +/- 2 s *versus* 107 +/- 25 s, Figure 3G). Cervical vagotomy eliminated responses to topical administration of substance P (Figure 3H; Movie S2). These observations indicate that topical administration of chemical substances to the pancreas elicits specific responses in peripheral axonal terminals. As shown below, the combination of two different stimulation approaches (intraductal and topical) together with proper controls made it possible to identify and study sensory mechanisms of vagal sensory transmission from the pancreas.

### Vagal afferent neurons innervating the pancreas are chemosensors

Using *in vivo* imaging of the nodose ganglion, we assessed the responsiveness of vagal sensory neurons to mechanical and chemical stimulation of the pancreas (Figure 4). Pancreata were stimulated mechanically either by light touch with forceps and gentle rinse of the exteriorized portion of the pancreas (mimicking the conditions of topical stimulus application) or by stretching the pancreas *via* intraductal injection of saline at a rate of 300 μl/min for 30-60 s (twice the pressure at which chemical stimuli were applied intraductally). In contrast to what has been described for sensory neurons innervating the stomach where nearly 70% of neurons respond to stretch (Williams et al., 2016; Zagorodnyuk et al., 2001), few neurons responded to mechanical distention of the pancreas (∼16%, 31/193, n = 3 mice; Figures 4B and 4C). The responses to mechanical stimuli were uncoordinated, had low amplitude, and were barely distinguishable from baseline neuronal activity. This lack of mechanical sensation is in line with the notion that the pancreas does not distend or contract physiologically.

**Figure 4.**
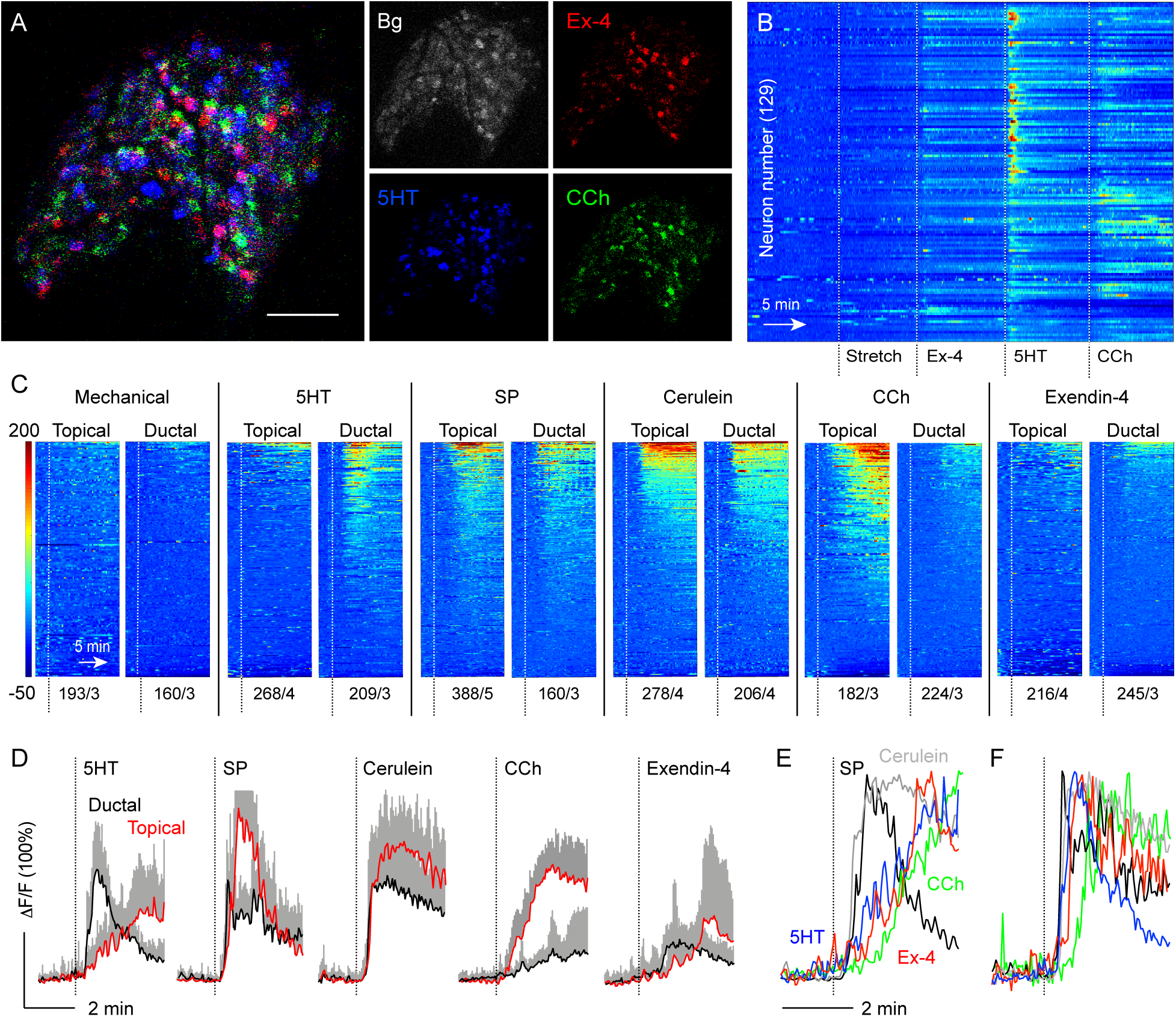
Pancreatic vagal afferents are chemosensors. (A) Spatial map of GCaMP6 fluorescence signals in the nodose ganglion at the peak of the response to intraductal application of exendin-4 (Ex-4, 10 μM, red), serotonin (5HT, 1 mM, blue), and carbachol (CCh, 100 μM, green). Background fluorescence is shown in gray. Scale bar, 100 μm. (B) Representative heatmap, showing color-coded dF/F Ca^2+^ responses of 129 single nodose ganglion neurons to pancreatic stimulation with indicated stimuli. Each row is a single cell, x-axis is time, color scale is dF/F (%), where fluorescence intensity increases from blue to red. (C) Series of heatmaps comparing nodose ganglion Ca^2+^ responses to intraductal and topical administration of mechanical and chemical stimuli to the pancreas. Number of neurons/number of animals are indicated under each heatmap. dF/F color scale also applies to (B). (D) Average traces of the top 15 responding neurons shown in (C). Ductal stimulation responders shown in black, topical stimulation responders shown in red. (E and F) Average traces as shown in (D), normalized to 100%, of responses to topical stimulation (E) or intraductal stimulation (F) of the pancreas with indicated stimuli.

We then explored the chemosensitive properties of pancreatic vagal afferents. We selected stimuli known to activate receptors on sensory axons directly (e.g. substance P and serotonin), activate pancreas tissues (e.g. carbachol), or both (e.g. cerulein and exendin-4). We tested a total of 15 molecules of which the aforementioned ones consistently elicited responses (Table S1). The route of stimulus administration (topical *versus* intraductal) affected the magnitude and incidence of responses to serotonin and carbachol (Figures 4C and 4D). Serotonin was more effective when applied intraductally, while carbachol was more effective when applied topically. Substance P, cerulein, and exendin-4, by contrast, elicited responses in a similar number of neurons regardless of the delivery route (Figure 4C).

Based on the response kinetics we could distinguish synchronized responses with steep slopes and non-synchronized responses with gradual slopes (Figures 4E and 4F, Table S2). Responses to substance P and cerulein were fast and had similar times to peak and slope coefficients, irrespective of the administration route. Carbachol, whether delivered topically or intraductally, elicited slow responses. The kinetics of the responses to serotonin and exendin-4 depended on the administration route. When applied intraductally, serotonin and exendin-4 elicited responses that were as fast as those to substance P and cerulein.

The numbers of responding neurons, the magnitude of the response, as well as the response kinetics suggest that serotonin, substance P, and cerulein directly activate their cognate receptors known to be expressed by vagal afferent neurons (5HT3R, Tac1, and Cckar, respectively; Kupari, et al., 2019). The slower kinetics of the responses to carbachol suggest an indirect effect, for instance through activation of the exocrine tissue which then stimulates sensory nerves. Serotonin stimulated sensory neurons much faster and more efficiently when applied intraductally, indicating that serotonin-responsive axons were located closer to the pancreatic duct. Of note, the majority of pancreatic islets are aligned along the major pancreatic ducts.

### Mouse pancreatic islets contain and release serotonin

Because vagal afferents in the pancreas were sensitive to serotonin, we searched for the endogenous source of serotonin that could activate these neurons. Although serotonin derived from enterochromaffin cells is released into the blood, most of it is stored in platelets and only a small fraction remains freely circulating (El-Merahbi et al., 2015). It is known that beta cells produce and secrete serotonin as a paracrine and autocrine signal (Almaça et al., 2016; K. Kim et al., 2015; Ohara-Imaizumi et al., 2013). Although serotonin levels are lower in mouse beta cells than in human beta cells (Almaça et al., 2016), basal beta-cell serotonin levels play a role in glucose stimulated insulin secretion, which is more evident under high fat diet-induced metabolic stress conditions (Kim et al., 2015). We therefore investigated if under normal physiological conditions, serotonin is produced and released from mouse pancreatic islets (Figure 5). Although variable, serotonin immunostaining in pancreatic islets was significantly higher than background levels in exocrine regions (Figures 5A and 5D, Figure S4). Serotonin was present primarily in the beta cells (Figures 5B and 5B’). No other region of the pancreas showed serotonin immunostaining (Figures 5A and 5D). We also detected the tryptophan hydroxylase-1 (Tph-1), the rate-limiting enzyme in serotonin synthesis, in islets (Figures 5C and 5C’).

**Figure 5.**
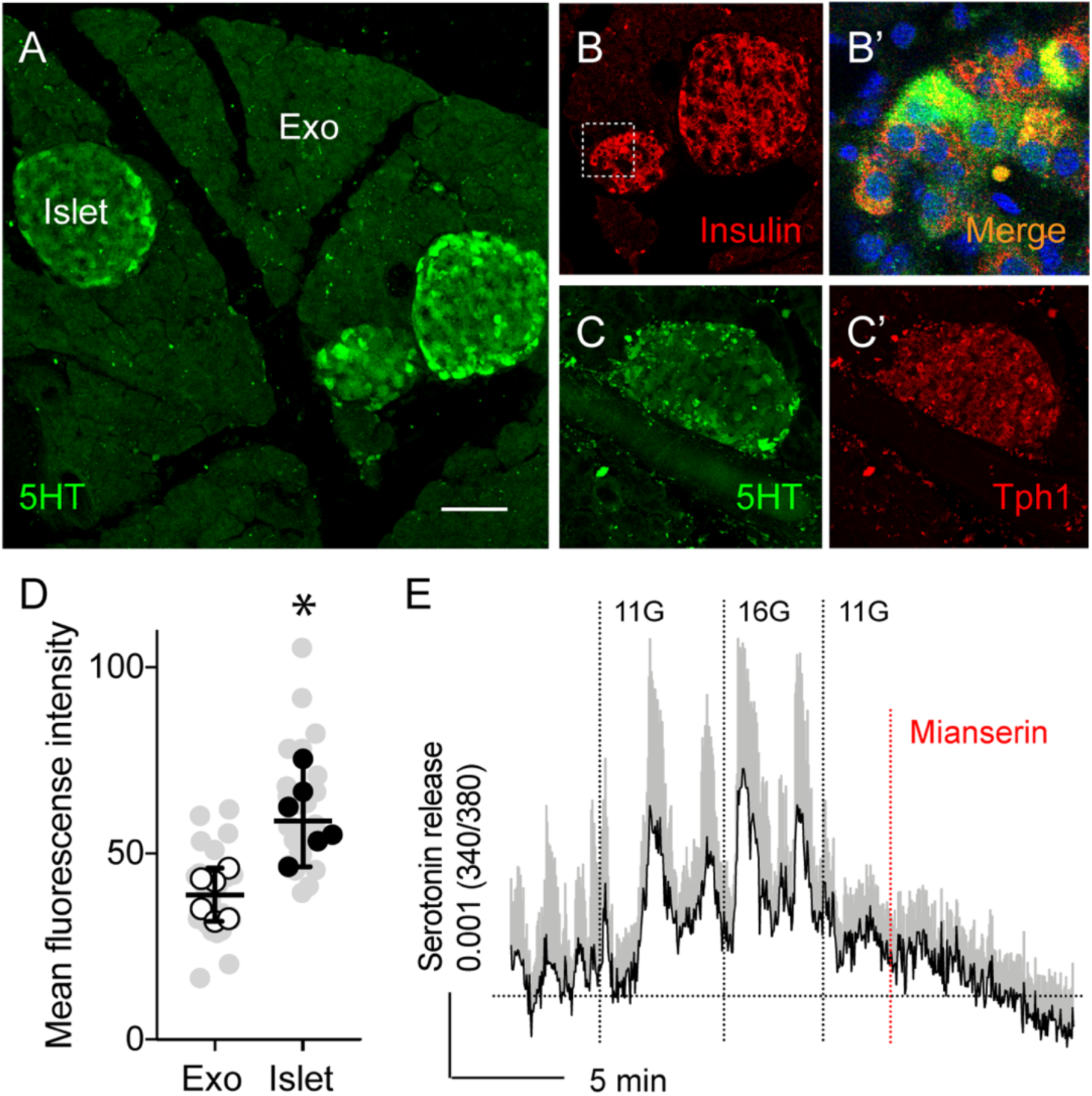
Mouse pancreatic islets contain and release serotonin. (A) Section of mouse pancreata showing that serotonin immunostaining (green) is limited to the islets and not detected in the exocrine tissue. Scale bar, 50 μm. (B) Section of mouse pancreata shown in A, contains beta cell insulin immunostaining (red). (B’) Enlarged view of the pancreatic islet shown in A and B, reflecting overlap in serotonin and insulin immunostaining (yellow). (C and C’) Section of mouse pancreata showing islet immunostained with serotonin (green, C) and Tph1 (red, C’). (D) Quantification of serotonin immunostaining as shown in (Figure S4). Immunostaining levels in endocrine regions (islet) were compared to those in exocrine regions (Exo) (n = 6 mice, 4-7 regions per animal; mean +/- SEM, paired t-test). Grey symbols represent values from all examined regions. (E) Stimulating mouse islets with increases in glucose concentration (3 mM to 11 mM to 16 mM) elicits serotonin secretion, as measured by biosensor cells (average of 9 cells +/- SEM). Responses are inhibited by the serotonin 5HT2C receptor antagonist mianserin (10 μM).

We used a biosensor cell approach to detect serotonin release from mouse islets (Almaça et al., 2016; Huang et al., 2005; Rodriguez-Diaz et al., 2012). We placed mouse islets on top of serotonin biosensor cells (CHO cells) expressing the serotonin receptor 2C (5-HT2C) loaded with the Ca^2+^ indicator fura-2. Stimulating islets with high glucose concentrations or with KCl depolarization elicited Ca^2+^ responses in biosensor cells (Figure 5E; Figure S5). These responses were blocked by the 5HT2 antagonist mianserin, confirming that the biosensor cells had detected serotonin. These data indicate that mouse pancreatic islets contain and release serotonin in a glucose-dependent manner.

### Serotonin activates axonal terminals in the pancreatic islet

The ionotropic 5HT3 receptor is a prominent serotonin receptor in vagal afferent neurons (Browning, 2015; Browning & Mendelowitz, 2003; Lacolley et al., 2006). We used transgenic mice in which the promoter for the 5HT3 receptor drives the expression of GFP (5HT3R-GFP mice; Chittajallu et al., 2013; Inta et al., 2008) to assess the presence of this receptor in sensory axons innervating the pancreatic islet. We found a high density of GFP-labeled fibers around and inside most examined islets (Figure 6A). These fibers also expressed the sensory axon marker CGRP (Figure 6A’ and 6A’’). To directly determine whether serotonin elicited responses in sensory fibers we adapted a living pancreas slice approach that enables studying the function of different tissue compartments in a preserved pancreas environment (Weitz et al., 2018). We prepared pancreas slices from Pirt-GCaMP3 mice and recorded activity of sensory fibers in the islet (Figures 6B-6E; Movie S3). We found that sensory fibers responded to stimulation of islet endocrine cells with high glucose concentration as well as to stimulation with serotonin (50 μM, Figures 6C-6E).

**Figure 6.**
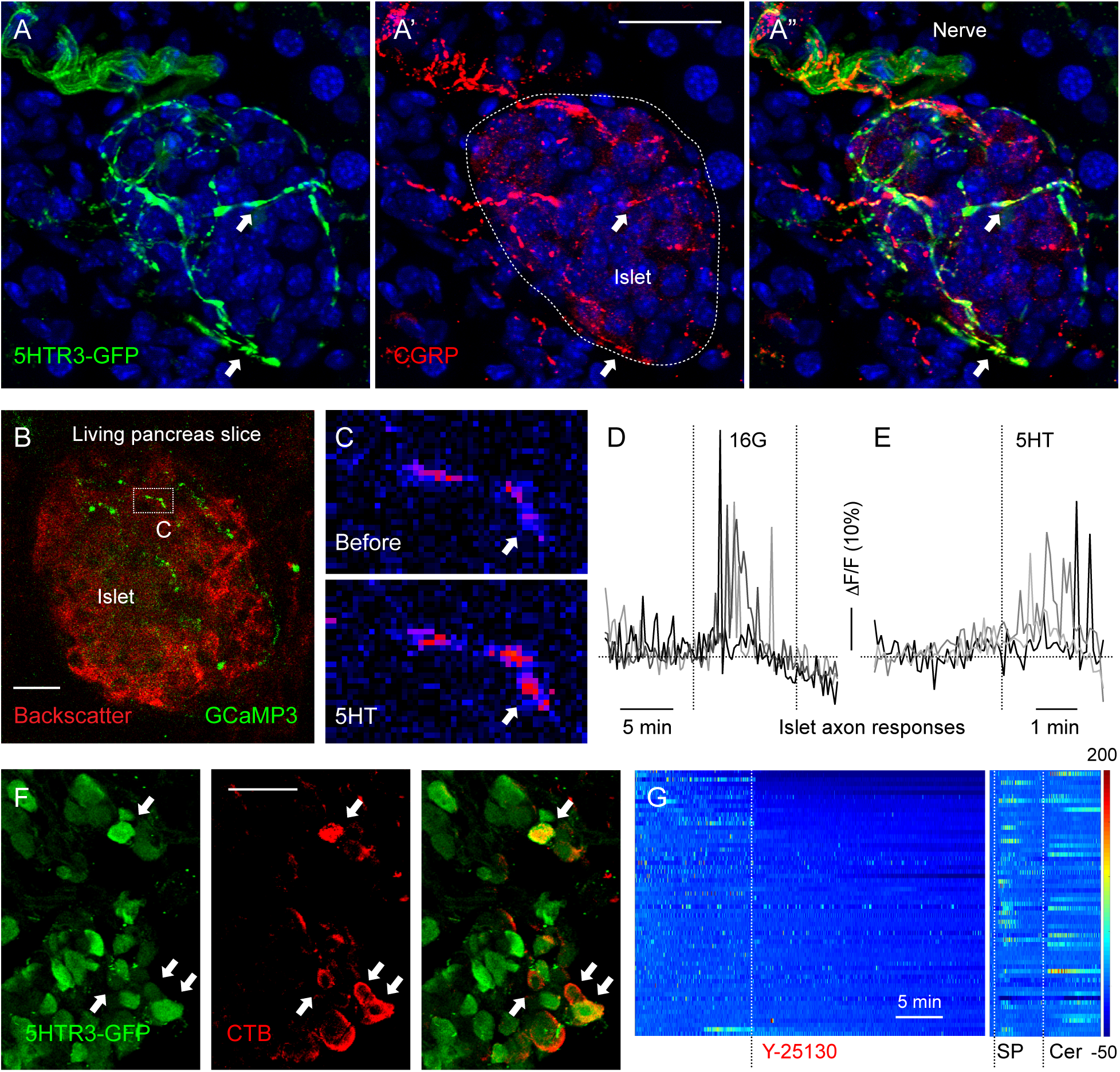
Serotonin activates axonal terminals in the mouse pancreatic islet. (A-A”) Pancreatic section from a 5HT3r-GFP reporter mouse showing an islet innervated by 5HT3r-expressing fibers that also immunostain for CGRP (red). Scale bar, 20 μm. (B) z-stack of confocal images of a living pancreatic slice from a Pirt-GCaMP3 mouse. Sensory fibers (green) can be seen in an islet visualized by backscatter (red). Scale bar, 20 μm. (C) Sequential images of sensory fiber shown in (B) displaying a Ca^2+^response to serotonin (5HT, 50 μM) perfused over the slice. Arrow points to a region showing an increase in GCaMP3 fluorescence (from blue to red in pseudocolor scale). (D and E) Representative traces of mean fluorescence intensity changes in the axonal terminals inside the pancreatic islet demonstrating that axonal terminals respond to an increase in glucose concentration (from 3 mM to 16 mM) and to 5HT (50 μM) stimulation. (F) Nodose ganglion section from a 5HT3R-GFP transgenic mice showing neurons retrogradely traced from the pancreas using CTB-594 tracer. Arrows point at traced neurons (red) that express the receptor (green). Scale bar, 100 μm. (G) Representative heatmap of changes in spontaneous activity of 61 nodose ganglion neurons in response to pancreatic topical application of the 5HT3r antagonist Y-25130 (1 mM; representative of 5 mice).

We confirmed that vagal afferent neurons that express 5HT3 receptors in the nodose ganglion project to the pancreas by retrograde CTB tracing in 5HT3R-GFP mice (Figure 6F). Using *in vivo* Ca^2+^ imaging of the nodose ganglion, we further determined that topical application to the pancreas of the 5HT3R antagonist Y-25130 (1 mM) diminished the spontaneous activity of a subpopulation of nodose neurons (∼30%), but did not affect responses to substance P or cerulein (Figure 6G). These findings not only show that vagal sensory neurons innervating the islet express 5HT3R and are able to sense serotonin directly, but also that their baseline activity partially originates from endogenous pancreatic serotonin signaling.

### Vagal sensory neurons respond to serotonin secreted from activated beta cells

We next assessed if vagal sensory neurons respond to serotonin derived from stimulated beta cells. Using *in vivo* Ca^2+^ imaging of the nodose ganglion, we recorded neuronal activity in response to topical application of tolbutamide to the pancreas (5 mM; Figure 7). Tolbutamide is a sulfonylurea that closes KATP channels, thus depolarizing beta cells and stimulating insulin secretion. Applying tolbutamide elicited small responses in a subpopulation of vagal sensory neurons (17%, n = 4 mice; Figures 7B, 7C, 7G, and 7H). Responses were delayed compared to those to substance P, suggesting that tolbutamide was acting indirectly *via* activation of beta cells. Topical application of the 5HT3R antagonist Y-25130 inhibited responses to tolbutamide, but not to cerulein (Figure 7H).

**Figure 7.**
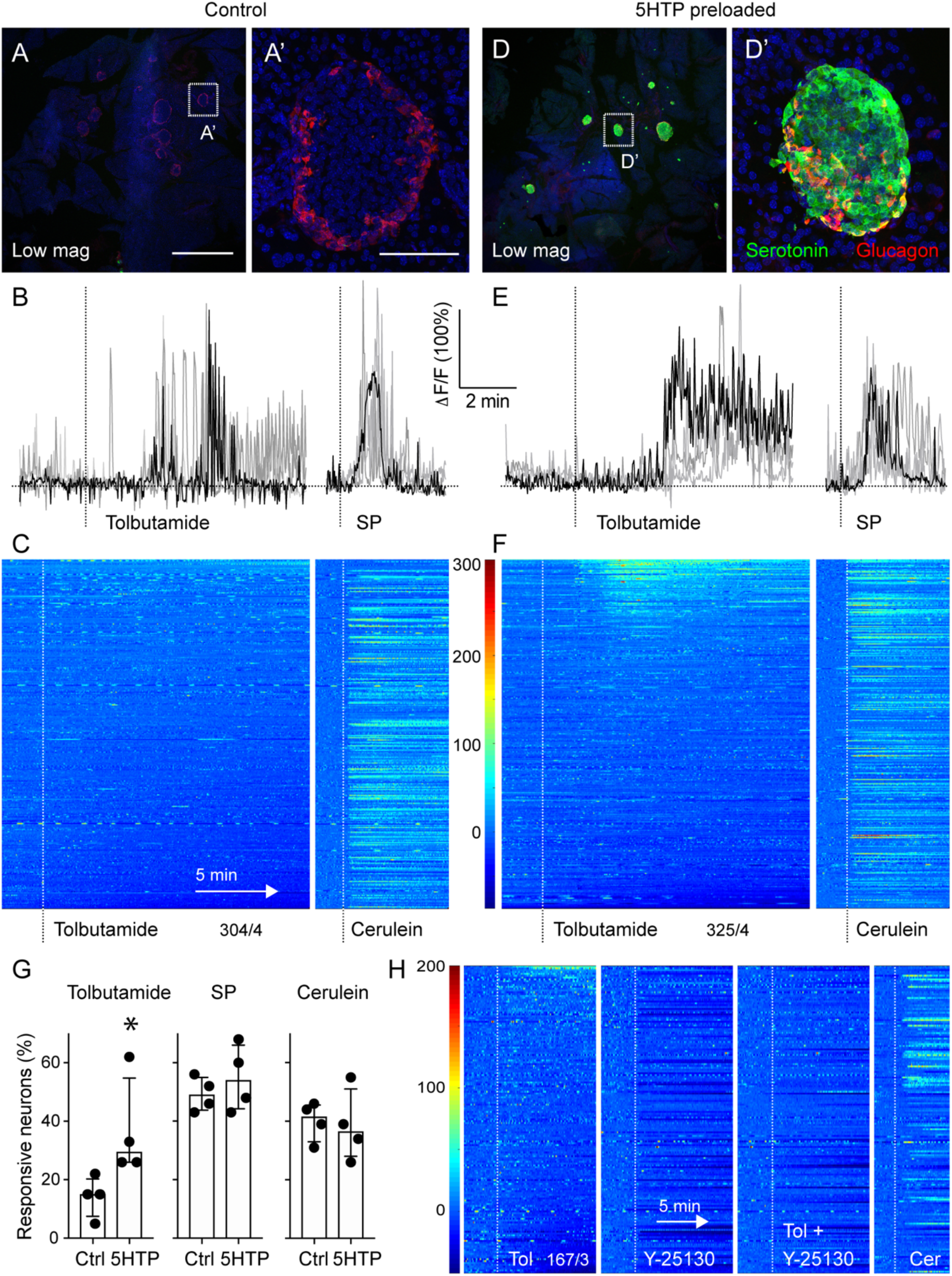
Vagal sensory neurons respond to serotonin secreted from beta cells. (A and D) z-stacks of confocal images of pancreatic sections from untreated mice (A) and mice preloaded with 5HTP (30 mg/kg; D). Serotonin immunostaining (green) is increased in islets (glucagon in red) from treated mice. (A’) and (D’) are higher magnifications of (A) and (D). Note that endogenous levels of serotonin in control tissue are not visible because imaging settings were adjusted for the increased serotonin levels in 5HTP-preloaded tissue. Scale bars, 1mm (A, D) and 50μM (A’, D’) (B and E) Representative traces of Ca^2+^ responses of nodose ganglion neurons to topical stimulation of the pancreas with tolbutamide (5 mM) and substance P (100 μM) in control (B) and 5HTP-preloaded (E) animals. (C and F) Heatmaps showing responses of nodose ganglion neurons to pancreatic stimulation with tolbutamide and cerulein in control mice (C, 304 neurons, n = 4 mice) and mice preloaded with 5HTP (F, 325 neurons, n = 4 mice). Each row is a single cell, x-axis is time, dF/F is color. (G) Quantification of the percentage of neurons responding to pancreatic stimulation in control and 5HTP preloaded animals (n = 4 for each group, Mann Whitney test, median +/- 95% CI). (H) Four sequential heatmaps reflecting changes in spontaneous activity and tolbutamide-evoked activity after administration of the 5HT3r antagonist Y25130 (1 mM). Notice that responses to cerulein persist.

We reasoned that the responses to tolbutamide were infrequent because islet endocrine cells represent a small fraction of the pancreatic mass (1-2%; Goodman, 2009). To amplify the effects of beta cell stimulation, we injected mice with the serotonin precursor 5-hydroxytryptophan (5HTP, 30 mg/kg, IP Zern et al., 1980). This treatment strongly and selectively increased serotonin levels in islets (Figures 7D and 7D’) but had only minor effects on glucose tolerance, as measured by intraperitoneal glucose tolerance test (12-24 hours after injection; Figure S6). In 5HTP-preloaded mice, the size and incidence of the responses to tolbutamide increased (>30% of vagal sensory neurons; Figures 7E-7G; Movies S4 and S5). 5HTP-preloading did not affect the magnitude and incidence of the responses to substance P and cerulein. These results demonstrate that beta cells use serotonin to communicate with vagal sensory neurons.

## DISCUSSION

Our study establishes that the pancreatic islet uses serotonin as a signaling molecule to communicate with the brain *via* vagal afferent neurons. This mechanism is supported by our findings showing that (a) sensory axons innervating the islet express serotonin receptors and respond to serotonin, (b) antagonizing serotonin receptors in the pancreas inhibits vagal sensory neuron responses to beta cell stimulation, and (c) increasing islet serotonin levels in beta cells amplifies neuronal responses to beta cell stimulation. By identifying serotonergic signaling between beta cells and sensory fibers as a peripheral transduction mechanism, our study reveals how beta cells activate vagal afferent neurons to make the brain aware of the process of insulin secretion.

Serotonin is an important signaling molecule in the islet. In the pancreas, serotonin is released from beta cells as a paracrine signal that helps control insulin and glucagon secretion (Almaça et al., 2016; Kim et al., 2015). The role of serotonin also includes intracellular serotonylation to regulate insulin exocytosis (Paulmann et al., 2009). Beta cells express all of the genes required to synthesize, package, and secrete serotonin (Ohta et al., 2011). In particular, the tryptophan hydroxylases TPH1 and TPH2, the two isoforms of the TPH enzyme that catalyze the rate limiting step in serotonin synthesis, are expressed in the mouse islet (Ohta et al., 2011). In line with these studies, we found that serotonin was produced and released from mouse islets. Our results indicate that regular levels of islet serotonin are physiologically relevant and sufficient to set the baseline activity in vagal afferent neurons. Activating sensory axons thus seems to be an important element in the functional repertoire of islet serotonin.

Serotonin signaling is a prominent transduction mechanism shared by vagal afferent axons throughout the viscera (Browning, 2015; El-Merahbi et al., 2015). The largest producers of serotonin in the body, the enterochromaffin cells of the gut, secrete serotonin in response to stimulation by nutrients and synaptically transmit this information to vagal afferent neurons by activating 5HT3 receptors (Bellono et al., 2017). Unlike enterochromaffin cells, whose massive release can increase serotonin levels systemically, beta cells secrete serotonin most likely for local consumption only. Given that the islet is the only source of serotonin in the pancreas, sensory axons need to be close to the islet if they are to detect serotonin. Indeed, we found that islets have the highest density of sensory axons in the pancreas. These axons are peripheral terminals of sensory neurons expressing 5HT3 receptors and respond to serotonin and activation of beta cells. Thus, by reaching into and around the islet, serotonin-sensitive axons put themselves in an ideal position to be exposed to the spatially restricted intra-islet serotonin.

It is important to emphasize that serotonin is packaged and released together with insulin in a glucose-dependent manner (Aspinwall et al., 1999; Ekholm et al., 1971). Serotonin has been used for decades in physiological experiments as a surrogate for insulin because serotonin is readily detected using electrochemical methods (e.g. amperometry; Barbosa et al., 1998; Bokvist et al., 2000; Aspinwall et al., 1999). By manipulating the serotonin content in insulin granules, we found that the magnitude of the afferent response depends on serotonin levels in beta cells. ATP, another molecule known to be released by beta cells (Hazama et al., 1998), did not stimulate nodose neurons. Insulin has been reported to stimulate dispersed nodose ganglion neurons and vagal afferents (Iwasaki et al., 2013; Niijima, 1981). Insulin receptor signaling is coupled to the PI3K/AKT pathway, barely affects the membrane potential, and is therefore difficult to reconcile with quick activation of neurons. Our *in vivo* studies show that vagal afferent responses to beta cell stimulation were dependent on serotonin signaling. We therefore propose that, in analogy to researchers who use amperometric measurements of serotonin to monitor insulin release in real time, afferent neurons detect serotonin as a proxy for insulin secretion. In our model, intra-islet serotonin changes the activity rate of sensory neurons in ways that reflect the secretory status of beta cells.

The production of serotonin in islets is remarkably variable. We found that even within the same pancreas islets had different serotonin levels. Under certain physiological conditions that involve changes in beta cell mass (e.g. pregnancy), serotonin synthesis increases tremendously, and serotonin serves as a paracrine signal that promotes beta cell proliferation (Kim et al., 2015; Ohara-Imaizumi et al., 2013). In the human islet, the levels of serotonin in beta cells also vary according to the BMI of the individual (Almaça et al., 2016). Moreover, some islets receive dense sensory supply, while others are innervated sparsely. All this heterogeneity suggests that not all islets contribute equally or constantly to islet-brain communication. We propose that islet serotonin released under normal conditions informs the brain about how much insulin is secreted. When serotonin levels are regulated in response to physiological challenges, however, serotonin may be a signal carrying information about functional adaptation and plasticity in the islet.

Recent studies are beginning to elucidate how vagal sensory neurons encode information originating from visceral organs (e.g. Bellono et al., 2017; Chang et al., 2015; Han et al., 2018; Kaelberer et al., 2018; Nonomura et al., 2016; Williams et al., 2016). A major challenge is that internal organs are very diverse in their functions and chemical composition, yet a single nerve collects sensory information from all of this diversity. It is still unclear if the vagus nerve is a mix of distinct, specifically tuned neuronal populations that possess unique strategies to sense the microenvironment in different organs (labeled lines) or if the diverse organs evolved similar and concerted transduction mechanisms across the vagus nerve (ensemble coding). Our study shows that vagal afferents monitor insulin secretion by detecting serotonin, using a mechanism that is similar to that used by the vagus nerve to respond to input from enterochromaffin cells (Bellono et al., 2017). Most neurons in the nodose ganglion express serotonin receptors (see Figure 6), supporting the notion that serotonin signaling is a common thread running through the physiological processes of digestion and energy homeostasis (Browning, 2015; Gershon & Tack, 2007). How sensory input from the islet affects responses in the central nervous system remains elusive. Our findings, combined with those of denervation studies, nevertheless suggest that without sensory input the autonomic centers of the brain may lose information about the functional status of the islet that is crucial for glucose homeostasis.

## ACKNOWLEDGEMENTS

This work was supported by the Diabetes Research Institute Foundation and National Institutes of Health grants R56DK084321 (A.C.), R01DK084321 (A.C.), R01DK111538 (A.C.), R01DK113093 (A.C.), U01DK120456 (A. C.) R33ES025673 (A.C.) and R21ES025673 (A.C.), K01DK111757 (J.A.), R21DK114418 (R.R.-D.), F31DK112596 (M.M.), the Leona M. and Harry B. Helmsley Charitable Trust grants G-2018PG-T1D034 and G-1912-03552, and by the American Heart Association 19POST34450054 (J.W.).

## AUTHOR CONTRIBUTIONS

M.M. conceived and designed the study, implemented all surgical approaches, developed protocols for the acquisition, analyses and presentation of data, collected anatomical and physiological data, and analyzed and interpreted the results. M.M. wrote the original draft of the manuscript. JW contributed the studies using living pancreas slices and its related data analyses. E.P. and A.T. contributed to the collection of the data including immunohistochemical experiments, IPGTTs, and data analyses. J.A. contributed to the conception of the study and to the serotonin detection studies. R.R.-D. contributed to the conception of the study, gave expert advice on immunohistochemical experiments, and performed the immunohistochemical studies to visualize sensory fibers in mouse and human pancreata. A.C. contributed to the conception and design of the study, data interpretation, reviewed the original draft of the manuscript, edited and approved the final version of the manuscript. A.C. is the guarantor of this work and has full access to the data and takes responsibility for the integrity of the work. All authors revised the original manuscript and approved the final version of the manuscript.

## DECLARATION OF INTERESTS

The authors declare no competing interests.

## EXPERIMENTAL PROCEDURES

### Mouse Models

For *in vivo* Ca^2+^ imaging experiments we used Snap25-GCaMP6s (The Jackson Laboratory, stock nr. 025111) and Pirt-Cre mice crossed to floxed GCaMP6s mice (The Jackson Laboratory, stock nr. 024106). For *ex vivo* Ca^2+^ imaging experiments in peripheral sensory terminals, we used Pirt-GCaMP3 knock-in mice. Pirt-Cre and Pirt-GCaMP3 knock-in mice were kindly donated by Dr. Stephan Roper, University of Miami (originally obtained from X. Dong, Johns Hopkins). It has been previously reported that in the these mouse models Pirt promoter drives expression of GCaMP in all sensory neurons (Shin et al., 2014; Wu et al., 2015). Snap25 promoter is reported to drive expression of GCaMP in most neuronal types throughout the brain (Madisen et al., 2015). We found that in these mouse models GCaMP was expressed in all neurons of vagal sensory ganglia. Peripheral nerve terminals, however, could only be seen in Pirt-GCaMP3 knock in mice. Only F1 heterozygous mice were used, from both sexes, 8-30 weeks old. We did not observe any gender or strain differences in our results.

For characterization of sensory fibers innervating the pancreas we used Htr3a-GFP mice [GENSAT, Tg(Htr3a-EGFP)DH30Gsat]. In these animals, GFP is expressed in cells that endogenously express the ionotropic serotonin receptor 5Ht3a. The pattern of transgene expression was previously validated by overlap with endogenous gene products by *in situ* hybridization and immunostaining (Chittajallu et al., 2013; Dvoryanchikov et al., n.d.; Inta et al., 2008). Breeder mice were kindly provided by Dr. Nirupa Chaudhari, University of Miami (originally obtained from Dr. Chris McBain, National Institutes of Health).

Experiments were conducted according to protocols and guidelines approved by the University of Miami Institutional Animal Care and Use Committee.

### Surgical exposure of the vagal sensory ganglion (nodose/petrosal/jugular complex) for imaging and injections

We used the same surgical approach, with slight adjustments, for *in vivo* Ca^2+^ imaging and for ganglion injections. In all procedures we exposed the left vagal ganglion. For *in vivo* Ca^2+^ imaging, mice were anesthetized with a combination of isoflurane (1-1.5%) and ketamine (60 mg/kg) and xylazine (5 mg/kg) throughout the surgery. Anesthesia levels were monitored throughout the surgery by hind paw withdrawal reflex. The combined anesthesia was necessary to maintain blood glycemia levels within the physiological range and to stabilize breathing. Anesthetized animals were placed in supine position on an infrared surgical warming pad (DCT-15, Kent Scientific) and the head was fixed in the head holder (SG-4N, Narishige). Tracheotomy was performed to facilitate respiration. The hypoglossal nerve was bluntly dissected and pulled to the lateral side with silk suture.

To access the ganglion, tissues were bluntly dissected using custom microdissection hooks. A first hook was placed on top of the trachea and common carotid artery at the level of the carotid bifurcation. This hook was secured to pull the tissues medially. A second hook was placed on top of the posterior belly of digastric muscle and the internal carotid and occipital arteries. This hook was secured to pull the tissues rostrally. A third hook was placed on top of the sternomastoid muscle and blood vessels supplying it. This hook was secured to pull the tissues laterally. Minimum tension was applied during retraction to minimize obstruction of blood flow. The animal was not allowed to recover from anesthesia and was euthanized by cervical dislocation at the end of the imaging session.

For tracer injections into the ganglion, the animal was anesthetized with 2-3% isoflurane throughout the surgery. The exposure of the ganglion was performed as described above, but without tracheotomy. Viral tracers were injected into the ganglion as previously described (Chang et al., 2015). Briefly, we used a pulled glass micropipette (30-80 μm diameter) connected to an automated nanoliter injector (2010, WPI) operated by a micromanipulator (MP-85, Sutter). The ganglion was pierced by the micropipette and 2-4 pulses of 69 nl tracer were injected into the ganglion. The wound was sutured, and animals were allowed to recover in a pre-warmed cage. Postoperatively, animals were given buprenorphen 5 μl/g subcutaneously, twice a day for three days, to minimize pain and discomfort. Animals were sacrificed for tissue collection 4 weeks post-surgery. Viral particles for anterograde neuronal tracing (AAV2 444-UbC-GFP titer 4.16×10^12^ and AAV8 733-mCherry titer 1.3×10^14^) were obtained from the Miami Project to Cure Paralysis Viral Core facility. Viral particles for retrograde neuronal tracing [pAAVrg-hSyn-DIO-hM3D(Gq)-mCherry titer 7×10¹² vg/mL were purchased from AddGene (stock number 44361)]. We observed higher transfection efficiency of the nodose neurons with the AAV8 serotype compared to the AAV2 serotype.

### Confocal imaging of the vagal sensory ganglion (nodose ganglion) *in vivo*

Anesthetized animals with the exposed left vagal ganglion were placed under a Leica TCS SP5 upright confocal microscope. Anesthesia levels were checked by the hind paw withdrawal reflex throughout the imaging session. We used a 10x/0.3NA dry objective with 11 mm working distance (#11506505, Leica) and resonant scanner for fast image acquisition. Because the anatomical position of the intact ganglion is not flat, we imaged in XYZT mode, spanning up to 500 μm in the z-plane to capture as much of the volume of the exposed ganglion as possible. The temporal resolution in all our experiments was 3 s at a digital resolution of 512×512 pixels. GCaMP fluorescence was recorded at 488Ex/510-550Em. The tissue did not dry out due to intact vascularization. We checked ganglion blood perfusion by injecting intravenously the mouse with DyLight 594 labeled lectin (DL-1067, Vector Labs, 594Ex/ 610-650Em) or 3kDa TRITC dextran (D3308, ThermoFisher, 568Ex/590-630Em).

### Infusion of the pancreas through the common bile duct

We used a previously described surgical approach of pancreas infusion for tracer delivery to the pancreas as well as for pancreas stimulation during *in vivo* Ca^2+^ imaging of the nodose ganglion (Guo et al., 2014). After abdominal midline incision, the common bile duct was exposed, and a small incision in the duodenum just below the ampulla of Vater was made using 31G needle. A 31G catheter (CMF31G, WPI) was inserted through the incision and through the ampulla of Vater into the common bile duct and clamped from the duodenal and hepatic sides using bulldog clamps. The catheter was connected to the 1 ml luer lock syringe controlled by an automated injector (55-2222, Harvard Apparatus).

For tracer delivery into the pancreas via intraductal infusion, the animal was anesthetized throughout the surgery with 2-3% isoflurane. The pancreas was infused with tracer (CTB, Fast Blue, AAV-retro) at a rate of 6 μl/min until a final volume of 7.5 μl/g of body weight was reached. After the infusion, the catheter was removed, muscle and skin were sutured, and the animal was allowed to recover in a pre-warmed cage. Postoperatively, animals were given buprenorphen 0.1 mg/kg subcutaneously, twice a day for three days, to minimize pain and discomfort. One-week post injection, we observed CTB and Fast Blue tracers throughout the pancreatic tissue but not in the duodenum. Altogether, these observations indicate that the tracer injection was specific. Although all three tracers provided similar tracing efficiencies at the level of the nodose ganglion, the AAV-retro was labeling the whole pathway from the pancreas to the brain stem.

For the *in vivo* imaging sessions, anesthesia was maintained as described in the “Surgical exposure of the vagal ganglion” section. To achieve pancreas distension, 300 μl of physiological buffer was infused at a rate of 300 μl/min. For chemical stimulation without pancreatic distension, 150 μl of stimulus was infused at a rate of 300 μl/min. Between stimuli, the pancreas was infused with physiological buffer at a rate of 5 μl/min for at least 5 minutes to wash out the remnants of the stimulus and to maintain the ductal tonus. The total volume infused into the pancreas per imaging session did not exceed 1 ml to eliminate pancreatic tissue damage and leakage of stimuli into the peritoneum. At the end of the imaging session, mice were not allowed to recover from anesthesia and were immediately euthanized by cervical dislocation.

### Pancreas exteriorization and topical stimulation

To exteriorize the pancreas, a small vertical incision was made on the left side of the mouse at the level of the spleen. Using a cotton swab, the tail of the pancreas and the spleen were gently exteriorized and placed on a 22 × 40 mm coverslip. The wound was sealed with silicone (KWIK-SIL, WPI) and a silicon border was built around the exteriorized portion of the pancreas, creating a chamber. It was essential to exteriorize the tissue very gently to avoid damage of the fine neuronal connections. The pancreas was submerged in physiological buffer inside the chamber. Solutions were added to the chamber with a pipette and removed by gentle aspiration.

### Preparation and imaging of living pancreatic slices

Acute pancreatic slices were prepared from Pirt-GCaMP3 transgenic mice for imaging of sensory axonal terminals, as previously described (Almaça et al., 2018; Marciniak et al., 2014; Weitz et al., 2018). After euthanasia, the abdomen was exposed and the pancreas infused through the common bile duct with 1.2% low gelling temperature agarose (39346-81-1, Sigma) dissolved in physiological buffer without BSA. After injection, the pancreas was extracted, cut into pieces, further embedded in agarose, and allowed to solidify at 4°C for 10 min. Pancreatic slices were cut on a vibratome (VT1000S, Leica) and incubated in physiological buffer (125 mM NaCl, 5.9 mM KCl, 2.56 mM CaCl2, 1 mM MgCl2, 25 mM HEPES, 0.1% BSA, pH 7.4) containing 3 mM glucose. Living pancreatic slices were placed in a perfusion imaging chamber (Warner Instruments) and imaged on a Leica TCS SP5 upright confocal microscope under continuous perfusion. GCaMP3 fluorescence was excited at 488 nm and emission detected at 510–550 nm. To identify pancreatic islets, we used the backscatter of 647 nm laser light. We recorded changes in GCaMP fluorescence induced by serotonin and KCl.

### Biosensor cells

Real time measurements of serotonin secretion were performed using fura-2 AM-loaded Chinese hamster ovary (CHO) cells expressing 5-HT2C receptors as previously described (Almaça et al., 2016; Huang et al., 2005; Rodriguez-Diaz et al., 2012).

### Intraperitoneal glucose tolerance test

Intraperitoneal glucose-tolerance tests (IPGTTs) were performed after overnight fasting. Mice were injected with 200–300 ml glucose solution (2 g/kg body weight) and blood glucose (IPGTT) was monitored at predetermined time points after the injection.

### Immunohistochemistry

Prior to tissue collection, animals were anesthetized with ketamine/xylazine (100/10 mg/kg, IP) and perfused transcardially with 4% PFA. Collected tissues (pancreas, duodenum, right and left vagal sensory ganglia, and the brain) were post fixed in 4% PFA overnight at 4°C, dehydrated in 30% sucrose overnight at 4°C, snap frozen and sectioned on a Leica CM-3050S cryostat.

Sensory ganglia were sectioned at 15-30 μm, while pancreas, brain and duodenum were sectioned at 40-50 μm. Sections were rinsed with 0.3% PBS-Triton X-100 and incubated in blocking solution (Universal blocking Reagent, Biogenex) in 0.3% PBS-Triton X-100. Afterwards, sections were incubated for 24 h at 20°C with primary antibodies diluted in blocking solution, rinsed and incubated with Alexa-Fluor conjugated secondary antibodies (1:500 in PBS) for 12-24 h at 20°C. Refer to the antibody table for the full list of primary and secondary antibodies. Slides were mounted with the VectaShield mounting medium (H-1000, Vector Laboratories). Confocal images of immunostained sections were acquired on an inverted Leica TCS SP5 confocal microscope.

### Data presentation, analyses and statistics

#### Quantification of sensory innervation

Immunohistochemistry images are presented as z-stack of confocal images, 8-10 images in a 40 μm section, z step = 4 μm. We used ImageJ software (https://imagej.nih.gov/ij/) to quantify the density of sensory innervation in the pancreatic islet and islet-surrounding exocrine tissue. For this quantification we used substance P immunostaining, which of all sensory markers tested provided the best signal to noise ratio with minimal background staining. We quantified mean gray value of substance P fluorescence in the maximal projections of confocal planes spanning the islet and surrounding exocrine tissue. Regions of interest in the islet were selected based on glucagon staining and dense DAPI staining. For the exocrine tissue, regions of interested were selected based on homogeneous DAPI staining of acini, avoiding endocrine, ductal, or blood vessel structures. Mean gray values of axon staining were calculated for several mouse strains (Figure S1). Average of mean gray values per animal are reported in Figure 1C.

#### Quantification of retrogradely traced neurons in the nodose ganglion

We used ImageJ software (https://imagej.nih.gov/ij/) to estimate the number of neurons labeled with the retrograde tracer. Data were calculated as percentage of nodose ganglion neurons stained with general neuronal marker NeuN in confocal planes.

#### Quantification of cytosolic Ca^2+^ Levels

To quantify changes in intracellular Ca^2+^ levels, we manually selected regions of interest (ROIs) around individual nodose ganglion neurons using the imageJ plugin Cell Magic Wand, developed by Fitzpatrick lab at the Max Planck Florida Institute. We only included neurons that expressed either stimulus-evoked or baseline Ca^2+^ responses. Thus, ROI selection was biased towards bright and responding neurons.

We measured changes in mean GCaMP fluorescence intensity using ImageJ. We used custom MatLab script for data analysis. Changes in fluorescence intensity were expressed as percentage changes over baseline (dF/F). The baseline was defined as the mean of the intensity values during the non-stimulatory control period of each recording. Most neurons exhibited Ca^2+^ signals during the non-stimulatory control period (baseline activity) as well as stimulus evoked responses. The strong baseline activity made it difficult to distinguish responders from non-responders based on amplitude and area under the curve, giving many false negatives and not adequately reflecting the data. For this reason, data were displayed as heatmaps including all recorded neurons. In the heatmaps, the response incidences, magnitudes and kinetics are visualized simultaneously. We further reported percentage of responding neurons based on inspection of individual Ca^2+^ traces by tree blinded independent observers.

To analyze response kinetics, we took the average of the 15 top responding neurons for each stimulus and normalized these averages to 100% (Figure 4E and 4F). Using GraphPad Prism, we then fitted a sigmoidal curve over each average trace. For each sigmoidal curve we calculated slope coefficient and time to peak (Table S2).

#### Data analyses

Sample sizes are indicated in main text, figure legends, or heatmaps (numbers in parentheses). Significance was determined by comparisons to the paired control group using paired t-test (Figures 1 and 5), multiple t-tests (Figure S6), or between indicated groups using a two-tailed Mann-Whitney test (Figure 7). All experiments involved biological, not technical replicates. We considered statistical significance when P values were lower than 0.05.

