## Supplementary Figures and tables for "Pancreatic Islets Communicate With the Brain *via* Vagal Sensory Neurons"

**SUPPLEMENTARY MATERIAL**

### Supplemental Figures

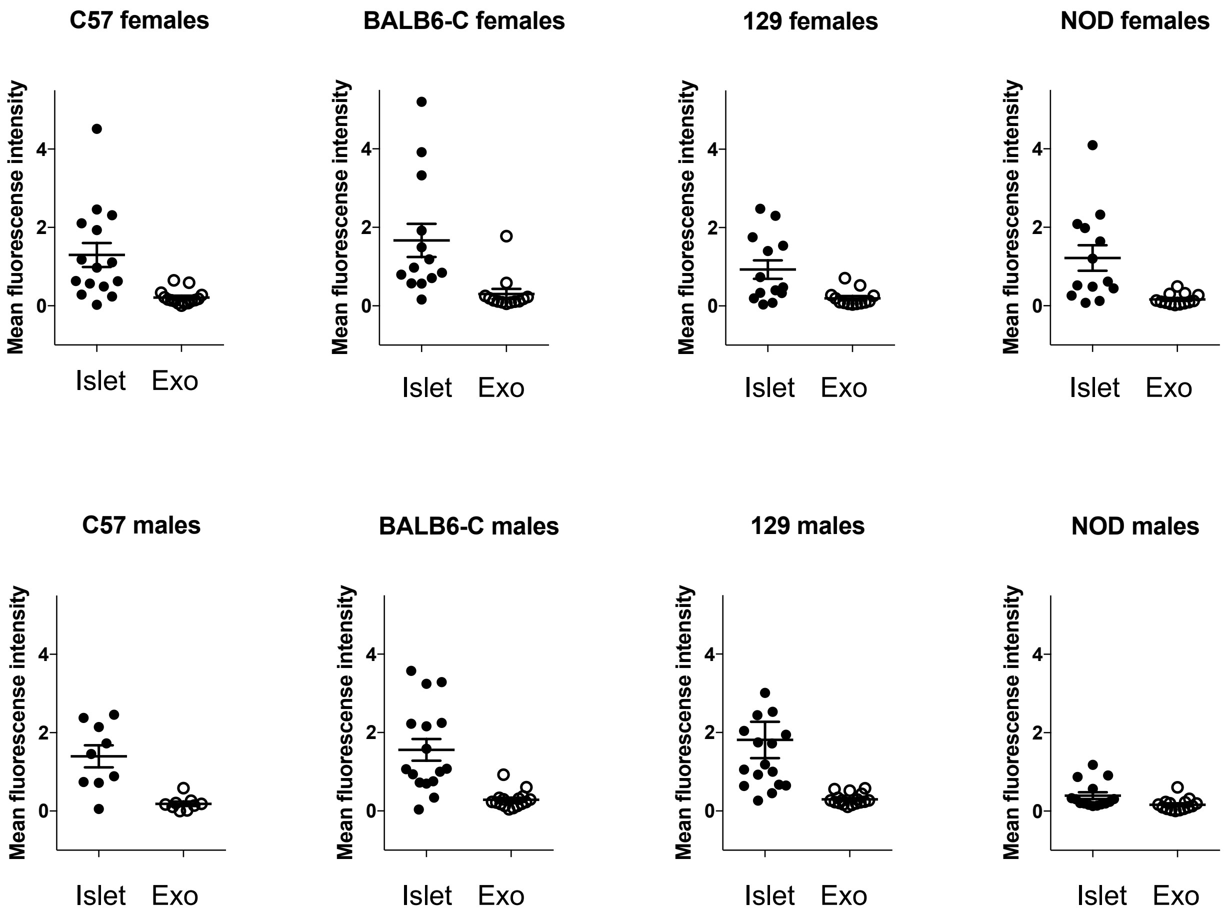


**Figure S1. Densities of sensory innervation in different mouse strains**
Quantification of the mean fluorescence intensity of substance P immunostaining in the islet and the surrounding exocrine tissue (n = 24 mice total, 3 mice for each category, 3-7 islets per mouse.

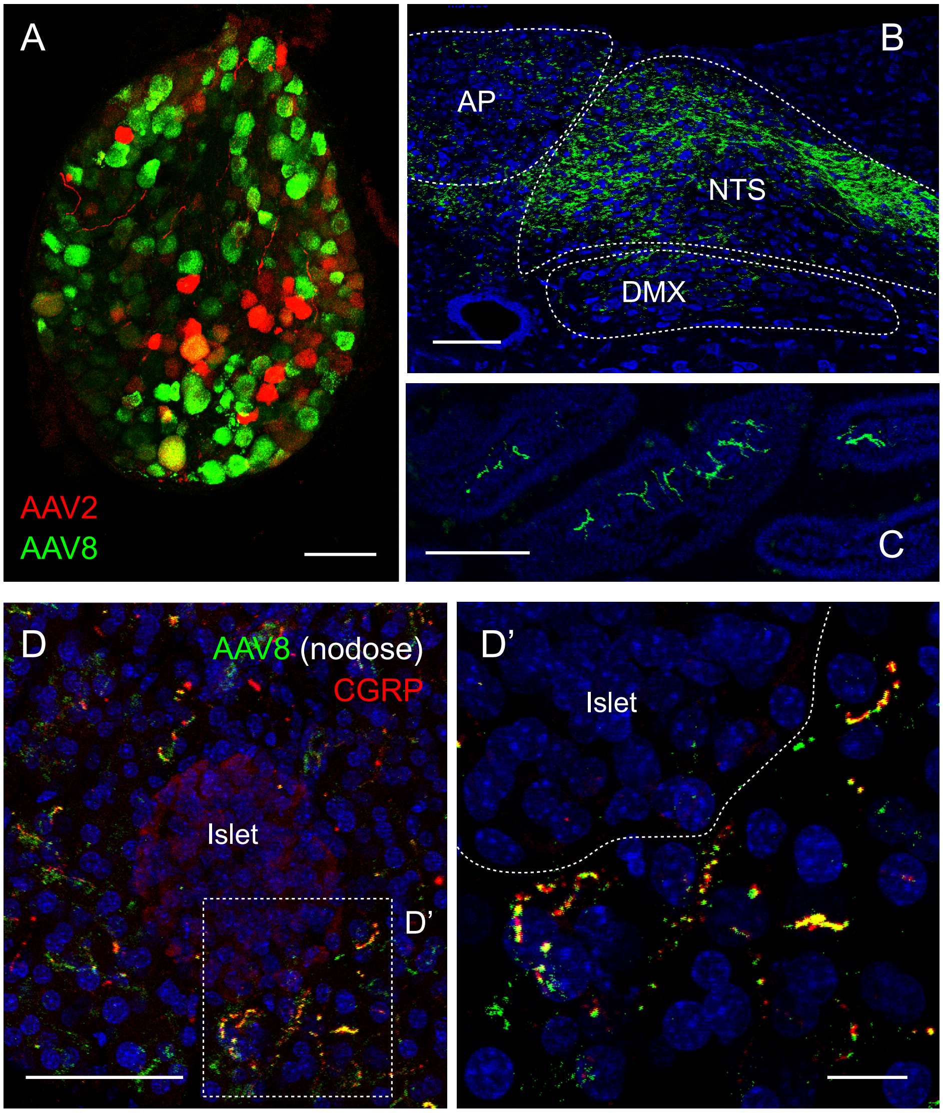


**Figure S2. Anterograde tracing from the nodose ganglion to the periphery and the brain**
(A) Section of the nodose ganglion with endogenous (non-amplified) expression of mCherry and GFP fluorescence 4 weeks after nodose ganglion transfection with AAV2 (red) and AAV8 (green). We observed better transfection and tracing efficiencies with AAV8 comparing to AAV2. Scale bar, 100 μm.
(B) Section of the caudal brainstem with endogenous (non-amplified) fluorescent protein expression in nerves traced from the nodose ganglion. Scale bar, 100 μm.
(C) Section of the duodenum, with immunohistochemical amplification of the fluorescence in the vagal afferents traced from the nodose ganglion (green). Scale bar, 100 μm.
(D and D') Section of the pancreas, with immunohistochemical amplification of the fluorescence in vagal afferents traced from the nodose ganglion(green), overlapping with CGRP staining (red). Scale bars, 10 μm.

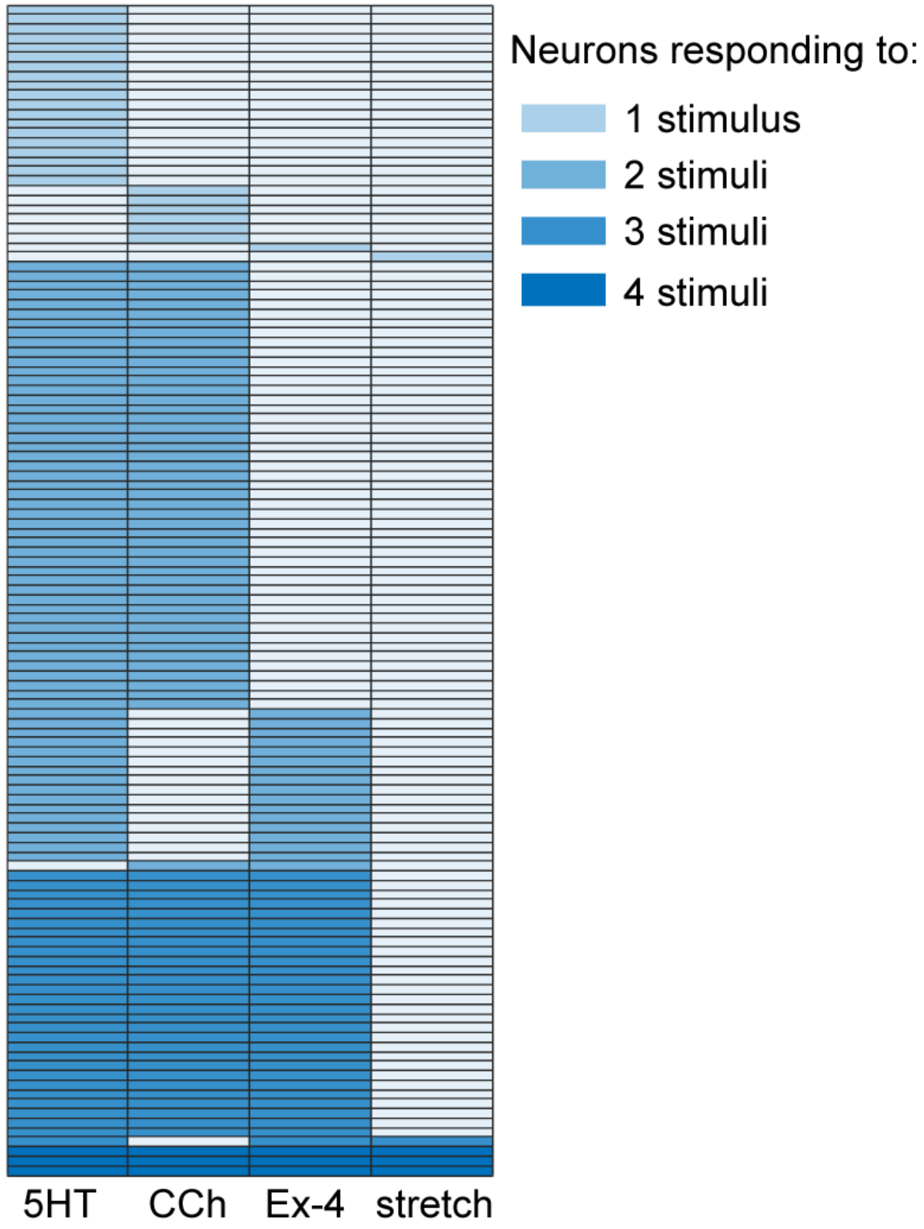


**Figure S3. Chemical specificity of vagal sensory neurons during intraductal stimulation of the pancreas**

Summary of the response profiles of 129 vagal afferents imaged in a single experiment as shown in Figure 4A and 4B. Each column indicates responses to a different compound specified at the bottom. Each row represents data from an individual nodose ganglion neuron. Most neurons responded to two stimuli: either 5HT and CCH or 5HT and Ex-4. Other neurons responded to one, three, and four stimuli.

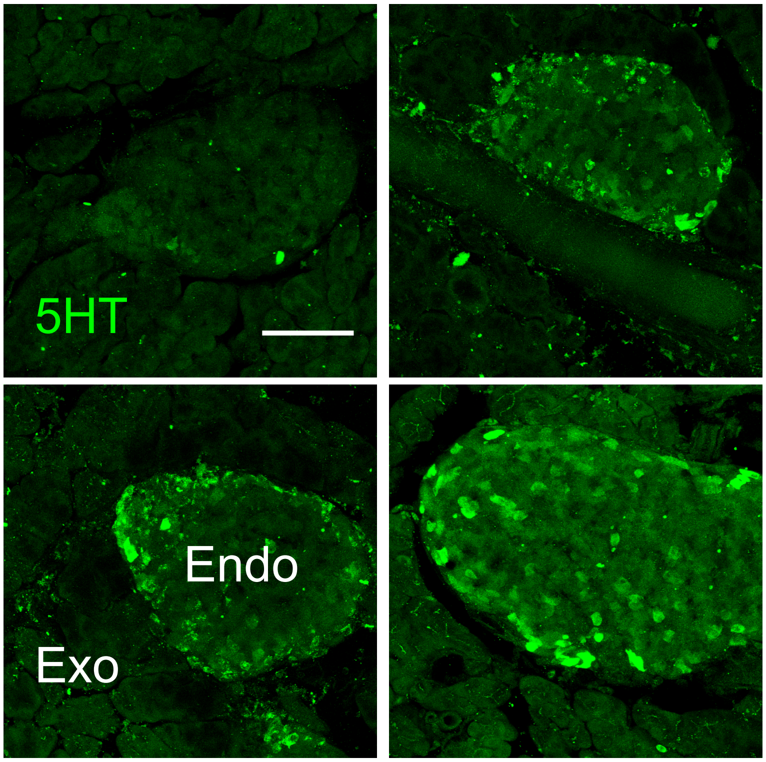


**Figure S4. Islets contain variable levels of serotonin**
Sections of mouse pancreata showing serotonin immunostaining (green) in different islets. Scale bar, 50 μm. For data quantification see Figure 5D.

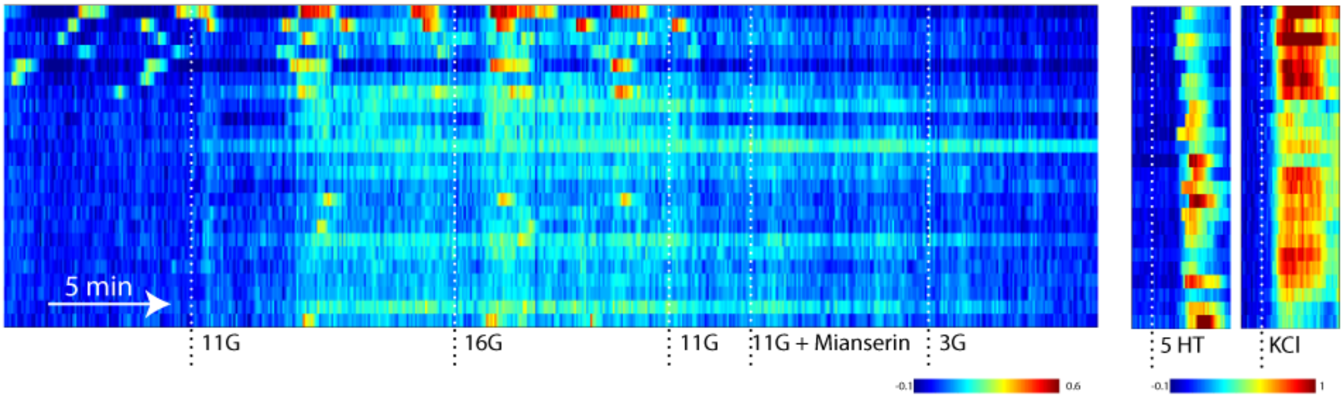


**Figure S5. Mouse pancreatic islets release serotonin**
Mouse islets were placed on a carpet of serotonin sensitive biosensor cells. Stimulating islets with increases in glucose concentration (3 mM to 11 mM to 16 mM) or with KCl depolarization (30 mM) elicited serotonin secretion, as measured by biosensor cells. Shown is a representative heatmap of fura-2 $Ca^{2+}$ responses of 24 single biosensor cells to islet stimulation. Responses were inhibited by the serotonin $5HT_{2C}$ receptor antagonist mianserin (10 µM). Serotonin (5HT, 10 nM) was added as a control. Each row is a single cell, x-axis is time, color scale is dF/F (%), where fluorescence intensity increases from blue to red.

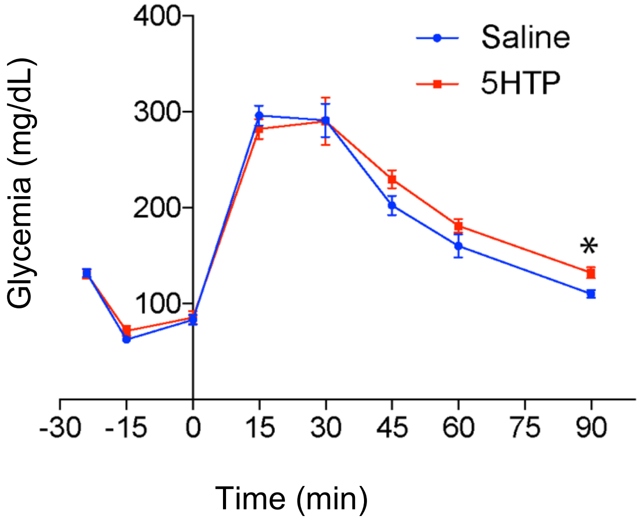


**Figure S6. Islet preloading with serotonin has a mild effect on glucose tolerance**
Intraperitoneal glucose tolerance test in control (blue) and serotonin-preloaded (red) mice (n = 7 mice per experimental group, mean +/- SEM, multiple t-tests, significant difference in glycemia is observed only at the 90 min readout). Glucose bolus was given at time 0.

| Stimuli | Concentration (μM) | Response strength |
|---|---|---|
| D-Glucose | 25000 | - |
| Tolbutamide | 1000 | + |
| 5HT | 1000 | ++ |
| ACh | 100 | + |
| CCh | 100 | ++++ |
| SP | 100 | ++++ |
| Kainate | 100 | + |
| ATP | 100 | - |
| Frosc/IBMX | 20/100 | +++ |
| Ex-4 | 10 | + |
| CGRP | 10 | - |
| Cyclosomatostatin | 10 | + |
| PP | 1 | + |
| Glucagon | 1 | + |
| Cerulein | 0.75 | +++ |

| | |
|---|---|
| - | < 10% |
| + | 10 - 20% |
| ++ | 20 - 30% |
| +++ | 30 - 40% |
| ++++ | > 40% |

**Table S1. List of chemical stimuli applied to the pancreas topically**

Fifteen compounds were tested at the indicated concentrations. Response strength was coded based on the percentage of nodose ganglion neurons responding to topical stimulation of the pancreas with the corresponding compound.

CCh = carbachol (cholinergic agonist)
SP = substance P
Frosc = forskolin
Ex-4 = exendin-4 (GLP-1 agonist)
PP = pancreatic polypeptide

| Stimuli | Slope coefficient | Time to peak |
| --- | --- | --- |
| 5HT_topical | 0.03471 | 165 |
| 5HT_ductal | 0.8665 | 63 |
| SP_topical | 0.3663 | 39 |
| SP_ductal | 1.386 | 33 |
| Cerulein_topical | 0.268 | 48 |
| Cerulein_ductal | 0.7401 | 48 |
| CCh_topical | 0.03705 | 210 |
| CCh_ductal | 0.1513 | 72 |
| Exendin-4_topical | 0.05732 | 144 |
| Exendin-4_ductal | 0.9477 | 66 |

**Table S2. Response kinetics of vagal sensory neurons stimulated from the pancreas**
Time to peak is given in seconds. Slope coefficient and time to peak were calculated as described in Methods.

**Supplemental movies**

**Movie S1. $Ca^{2+}$ responses of vagal afferent neurons to stimulation of the pancreas *in vivo***
Movie composed of a series of maximal projections of confocal images taken every 4.5 s of the left nodose ganglion of a Pirt-GCaMP6 mouse. Serotonin, substance P, and cerulein were applied sequentially to the exteriorized pancreas (related to Figure 3).

**Movie S2. Cervical vagotomy eliminates responses to pancreas stimulation**
Movie composed of a series of maximal projections of confocal images taken every 3 s of the left nodose ganglion of a Pirt-GCaMP6 mouse. Substance P was applied before and after vagotomy (related to Figure 3). Neurons were stimulated at the end of the experiment with KCl applied to the cut trunk of the nerve.

**Movie S3. $Ca^{2+}$ responses of sensory fibers in living pancreas slices**
Movie composed of a series of maximal projections of confocal images taken every 7.4 s of sensory fibers in an islet in a pancreas slice from a Pirt-GCaMP3 mouse. Serotonin (50 μM) was applied to the pancreas slice at the indicated time (related to Figure 6). Islet endocrine cells are visualized by their strong backscatter signal.

**Movie S4. $Ca^{2+}$ responses of vagal afferent neurons to stimulation of beta cells *in vivo***
Movie composed of a series of maximal projections of confocal images taken every 2 s of the left nodose ganglion of a Pirt-GCaMP6 mouse. Serotonin, the beta cell stimulus tolbutamide, substance P, cerulein, and ATP were applied sequentially to the exteriorized pancreas (related to Figure 7).

**Movie S5. $Ca^{2+}$ responses of vagal afferent neurons to stimulation of beta cells *in vivo* in a mouse treated with 5HTP**
Movie composed of a series of maximal projections of confocal images taken every 3 s of the left nodose ganglion of a pirt-GCaMP6 mouse. Serotonin, the beta cell stimulus tolbutamide, substance P, and cerulein were applied sequentially to the exteriorized pancreas (related to Figure 7).
